# Frequency of mutations in 21 hereditary breast and ovarian cancer susceptibility genes among high-risk Chinese individuals

**DOI:** 10.1101/514539

**Authors:** Di Shao, Shaomin Cheng, Fengming Guo, Yuying Yuan, Kunling Hu, Zhe Wang, Xuan Meng, Xin Jin, Yun Xiong, Xianghua Chai, Hong Li, Yu Zhang, Hongyun Zhang, Jihong Liu, Mingzhi Ye

## Abstract

To determine the prevalence and clinical prediction factors associated with deleterious mutations among 882 high-risk Chinese individuals who underwent multigene panel testing for hereditary breast and ovarian cancer (HBOC) risk assessment. Subjects were selected from individuals referred for genetic testing using a 21-gene panel (Oseq-BRCA) between January 2015 and March 2018. The distribution and prevalence of deleterious mutations were analyzed for the full cohort as well as subtypes. Overall, 176 deleterious mutations were observed in 19.50% (n = 172) individuals. Of these, 26 mutations are not reported in public databases and literatures. In the ovarian cancer only subgroup, 115 deleterious mutations were identified in 429 patients (48.6%). Patients with ovarian cancer with mutations were enriched for a family history of breast or ovarian cancers (p < 0.05). In the breast cancer only subgroup, 31 deleterious mutations were identified in 261 patients. Most mutations occurred in *BRCA1* (8; 25.8%) and *BRCA2* (11; 35.5%). An additional 12 deleterious mutations (38.7%) were found in 7 other susceptibility genes. An increased frequency of mutation rate (57.9%) was observed in the subgroup of subjects with histories of both breast and ovarian cancer. Taken together, 19.50% of individuals carried a deleterious mutation in HBOC susceptibility genes in our cohort. Subgroup of subjects with histories of both breast and ovarian cancer had the highest prevalence of mutations. Our results highlighted the genetic heterogeneity of HBOC and the efficiency of multigene panel in performing risk assessment.

## Introduction

Breast cancer is one of the most common malignant tumors in females in China. ^1^ Although ovarian cancer is less common than breast cancer, its mortality is high.^2^ Inherited mutations of *BRCA1* and *BRCA2* are pathogenetic in a majority of HBOC patients.^3,4^ In addition to *BRCA1* and *BRCA2*, studies have confirmed associations with other genes such as *ATM, BRIP1, CHEK2, RAD50, RAD51C*, and *TP53*.^5-8^ Moreover, it has been indicated that *TP53, PTEN, STK11*, and *CDH1* mutation carriers who may suffer from Li-Fraumeni syndrome, Cowden syndrome, Peutz–Jeghers syndrome, and hereditary diffuse gastric cancer syndrome, respectively, have high risks of breast cancer.^9-13^

In consideration of associations between genes and disease, the National Comprehensive Cancer Network (NCCN) recommends genetic testing of 19 genes in breast and ovarian cancer patients.^14^ Timely and effective genetic testing could provide professional counseling and clinical management for patients and at-risk relatives. As *BRCA1/2* carriers display the high sensitivity to inhibitors of poly-ADP-ribose polymerase (PARP), genetic testing could also help patients in choosing therapies.^15,16^

The traditional genetic testing methods, such as polymerase chain reaction (PCR) based assay, and denaturing gradient gel electrophoresis (DGGE) mutation scanning system have been applied to detect *BRCA1/2* widely in China.^8,17^ However, *BRCA1/2* are high-risk tumor suppressor genes without significant mutation hotspots; as a result, some mutations would be missed by conventional approaches. Results from recent studies confirmed that NGS showed multiple advantages in cancer genetic testing in terms of time and cost effectiveness.^18-20^ However, there is insufficient related reports on HBOC patients of the Chinese population.

To investigate the mutation frequency among individuals with a suspected HBOC risk in Chinese population, we used multi-gene testing to reveal the distribution and prevalence of deleterious germline mutations among 882 patients with a suspected HBOC risk in 21 HBOC heredity susceptive genes. Our results evaluated the benefits and limitations of multi-gene panel testing and provided insights of choosing appropriate multi-gene tests to assess the risk of hereditary cancer.

## Materials and Methods

### Participants

Subjects were selected from patients referred for genetic testing using a 21 gene panel Oseq-BRCA (BGI Genomics, Shenzhen, China) between January 2015 and March 2018. The enrollment criteria of this study were based on the current NCCN for genetic risk evaluation for HBOC: (1) diagnosed at any age with ovarian cancer or pancreatic cancer; (2) diagnosed with breast cancer with one or more of the following: diagnosed ≤ 50y, diagnosed with triple-negative breast cancer ≤ 60y, two or more separate breast cancer primaries, breast cancer at any age with at least one close blood relative with: breast cancer age ≤ 50y, male breast cancer, pancreatic cancer or ovarian cancer, and breast cancer at any age with at least two close blood relatives with breast cancer; (3) had a first- or second-degree relative with one or more of the following: breast cancer diagnosed ≤ 45y, ovarian cancer, male breast cancer or pancreatic cancer. (4) three or more close blood relatives on the same side of the family diagnosed with any cancer (14). Demographic and clinical information, including gender, personal cancer history, and family cancer history, were collected from test requisition forms (TRFs) by ordering clinicians at the time of testing. All patients signed informed consents approved by the Institutional Review Boards of BGI Genomics.

### NGS library construction and gene capture

Genomic DNA (gDNA) was extracted from participants’ peripheral blood samples using the Qiagen Blood Midi Kit (Qiagen, Hilden, Germany). DNA concentration and quality were assessed by Qubit (Life Technologies, Gaithersburg, MD, USA) and agarose gel electrophoresis. Genomic DNA (250 ng) was used for the sequencing library construction. Briefly, the gDNA was fragmented randomly by the Covaris LE220 sonicator (Woburn, MA) to generate gDNA fragments with a peak of 250 bp and then subjected to three enzymatic steps: end-repair, A-tailing, and adapter ligation. DNA libraries were purified with Agencourt Ampure XP beads (Beckman-Coulter, Indiana, USA), and PCR was performed during which a unique 8 bp barcode was added to label each sample. Five to ten PCR products were pooled equally and hybridized to a custom hereditary cancer panel (Roche NimbleGen, Madison, USA). Hybridization product was subsequently purified, amplified, and qualified. Finally, sequencing was performed with paired end and barcode on the BGISEQ-500 sequencer or Hiseq 2000 (Illumina, San Diego, CA) following the manufacturer’s protocols.

### Sequencing data analysis and mutation calling

Raw fastq data generated by the sequencer was first filtered by SOAPnuke to exclude low quality reads. The clean reads were then aligned to the reference human genome (UCSC hg19) using the BWA ALN algorithm. Single nucleotide variants (SNVs) were detected by the Genome Analysis Toolkit (GATK) Unified Genotyper. Small insertions and deletions (InDels) were called using GATK Haplotype. Copy number variants (CNVs) were called using read-depth analysis. All above variants were further filtered by quality depth, strand bias, mapping quality, and reads position. Finally, each variant was annotated with respect to gene location and predicted function in Human Genome Variation Society (HGVS) nomenclature and was ready prepared for interpretation.

### Data interpretation

Interpretation was focused on variants in 21 selected susceptibility genes in HBOC (Table 1). These 21 genes were selected through NCCN guidelines and published research articles, and they include core genes in the Fanconi anaemia (FA) pathway and HR genes (14). Variants were classified into the following 5 categories according to the American College of Medical Genetics (ACMG) recommendations: class 1, benign; class 2, likely benign; class 3, variant of uncertain significance (VUS); class 4, likely pathogenic(LP); and class 5, pathogenic (P).^21^ Population allele frequencies were collected from NCBI dbSNP, HapMap, 1000 human genome dataset, and an internal database of 100 Chinese healthy adults. Individuals with likely pathogenic or pathogenic variants were defined as having deleterious variants. Every deleterious variant was validated by qPCR, Sanger sequencing, or time of flight mass spectrometry.

**Table 1.** List of the 21 tested genes.

| **Risk Category** | **Genes** | **Gene names** |
| --- | --- | --- |
| *BRCA1/2* | 2 | *BRCA1, BRCA2* |
| *BRCA Pathway/Moderate Risk* | 9 | *ATM, BARD1, BRIP1, CHEK2, MRE11A, NBN, PALB2, RAD50, RAD51C* |
| *High Penetrant* | 4 | *CDH1, PTEN, STK11, TP53* |
| *Lynch Syndrome* | 5 | *MLH1, MSH2, MSH6, PMS1, PMS2* |
| *Other Moderate Risk gene* | 1 | *MUTYH* |

## Results

### Participant Characteristics

A total of 1,175 individuals were referred to our clinical test center for Oseq-BRCA multigene testing, from which 882 participants were included in our study based on NCCN guideline. Demographics for these 882 subjects are shown in Table 2. Among them, 709 samples had a diagnosis of either breast cancer or ovarian cancer, while 173 additional samples had a family history of cancer. Of participants with a cancer diagnosis, 261 subjects with a personal history of breast cancer, 429 subjects had a personal history of ovarian cancer, and 19 had personal histories of both breast and ovarian cancer. Age at diagnosis ranged from 13 to 80, with an average age of 47 years old. Among 173 subjects who did not have a cancer diagnosis, 108 (12.2%) had at least one first- or second-degree relative with breast cancer only, and 96 (10.9%) had a relative with ovarian cancer only.

**Table 2.** Demography and clinical characteristics.

|  | **Total**<br>**(n = 882)** | **BC**<br>**(n = 261)** | **OC**<br>**(n = 429)** | **BC & OC**<br>**(n = 19)** | **FHx**<br>**(n = 173)** |
| --- | --- | --- | --- | --- | --- |
| **Sex** |  |  |  |  |  |
| **Male** | 29 (3.3%) | - | - | - | 29 (16.8%) |
| **Female** | 853 (96.7%) | 261 (100%) | 429 (100%) | 19 (100%) | 144 (83.2%) |
| **Age at testing (yr)** |  |  |  |  |  |
| **[-,35]** | 170 | 78 | 22 | 0 | 70 |
| **(35,50)** | 369 | 165 | 130 | 5 | 69 |
| **[50, +]** | 340 | 17 | 276 | 14 | 33 |
| **NA** | 3 | 1 | 1 | 0 | 1 |
| **Mean(±SD)** | 47.0(±12) | 39.8(±7) | 53.5(±10) | 57.1(±10) | 40.4(±12) |
| **Median** | 47 | 40 | 54 | 57 | 38.5 |
| **Range** | 13-80 | 20-62 | 24-79 | 39-80 | 13-78 |
| **Family history** |  |  |  |  |  |
| **BC** | 108 | 30 | 31 | 0 | 47 |
| **OC** | 96 | 3 | 25 | 2 | 66 |
| **BC & OC** | 21 | 2 | 3 | 0 | 16 |

Abbreviations:
BC, Breast cancer; OC, Ovarian cancer; BC & OC, Breast and ovarian cancer; FHx, subjects recruited based on family cancer history; SD, standard deviation.

### Deleterious mutations identified in this cohort

Exons and splice sites of 21 HBOC susceptibility genes were examined for mutations by Oseq-BRCA in all 882 recruited participates. Overall, 176 deleterious (LP/P) mutations were observed in 19.50% (n = 172) individuals (Table 3). Of these mutations, 89 (50.6%) were found in *BRCA1*, 49 (27.8%) in *BRCA2*, and 38 (21.6%) mutations in 14 other susceptibility genes (Figure 1A, Figure 2). In addition, 2 individuals with ovarian cancer carried mutations in both *BRCA1* and another gene (*TP53* or *MRE11A*). Additionally, 2 individuals with breast cancer had mutations in both *CHEK2* and another gene (*BRCA2* or *TP53*). Deleterious mutations were identified in all individual genes, except *ATM, PTEN, CDH1, BARD*1 and *PMS2*.

**Table 3.** Frequency of mutations by personal cancer history.

| Gene | No. of Individuals (%) |  |  |  |  |
| --- | --- | --- | --- | --- | --- |
|  | Total<br>(n =882) | BC<br>(n =261) | OC<br>(n =429) | BC & OC<br>(n =19) | FHx<br>(n =173) |
| **BRCA1/2** |  |  |  |  |  |
| *BRCA1* | 89 (10.09)<sup>cd</sup> | 8 (3.07) | 66 (15.38)<sup>cd</sup> | 10 (52.63) | 5 (2.89) |
| *BRCA2* | 49 (5.56)<sup>a</sup> | 11 (4.21)<sup>a</sup> | 33 (7.69) | - | 5 (2.89) |
| **BRCA Pathway/Moderate Risk** |  |  |  |  |  |
| *BRIP1* | 9 (1.02) | 2 (0.77) | 5 (1.17) | - | 2 (1.16) |
| *CHEK2* | 7 (0.79)<sup>ab</sup> | 5 (1.92)<sup>ab</sup> | 1 (0.23) | - | 1 (0.58) |
| *MRE11A* | 3 (0.34)<sup>c</sup> | 1 (0.38) | 1 (0.23)<sup>c</sup> | - | 1 (0.58) |
| *NBN* | 1 (0.11) | - | - | - | 1 (0.58) |
| *PALB2* | 1 (0.11) | 1 (0.38) | - | - | - |
| *RAD50* | 1 (0.11) | - | 1 (0.23) | - | - |
| *RAD51C* | 4 (0.45) | 1 (0.38) | 2 (0.47) | - | 1 (0.58) |
| **High Penetrant** |  |  |  |  |  |
| *STK11* | 1 (0.11) | - | 1 (0.23) | - | - |
| *TP53* | 2 (0.23)<sup>bd</sup> | 1 (0.38)<sup>b</sup> | 1 (0.23)<sup>d</sup> | - | - |
| **Lynch Syndrome** |  |  |  |  |  |
| *MLH1* | 2 (0.23) | - | - | - | 2 (1.16) |
| *MSH2* | 3 (0.34) | - | 3 (0.70) | - | - |
| *MSH6* | 1 (0.11) | - | 1 (0.23) | - | - |
| *PMS1* | 1 (0.11) | 1 (0.38) | - | - | - |
| **Other Moderate Risk gene** |  |  |  |  |  |
| *MUTYH* | 2 (0.23) | - | - | 1 (5.26) | 1 (0.58) |
| **TOTAL** | **176(19.95)** | **31(11.88)** | **115(26.81)** | **11(57.89)** | **19(10.98)** |

<sup>a</sup>one subject had both *BRCA2* and *CHEK2* mutation
<sup>b</sup>one subject had both *CHEK2* and *TP53* mutation
<sup>c</sup>one subject had both *BRCA1* and *MRE11A* mutation
<sup>d</sup>one subject had both *BRCA1* and *TP53* mutation

**Figure 1.**
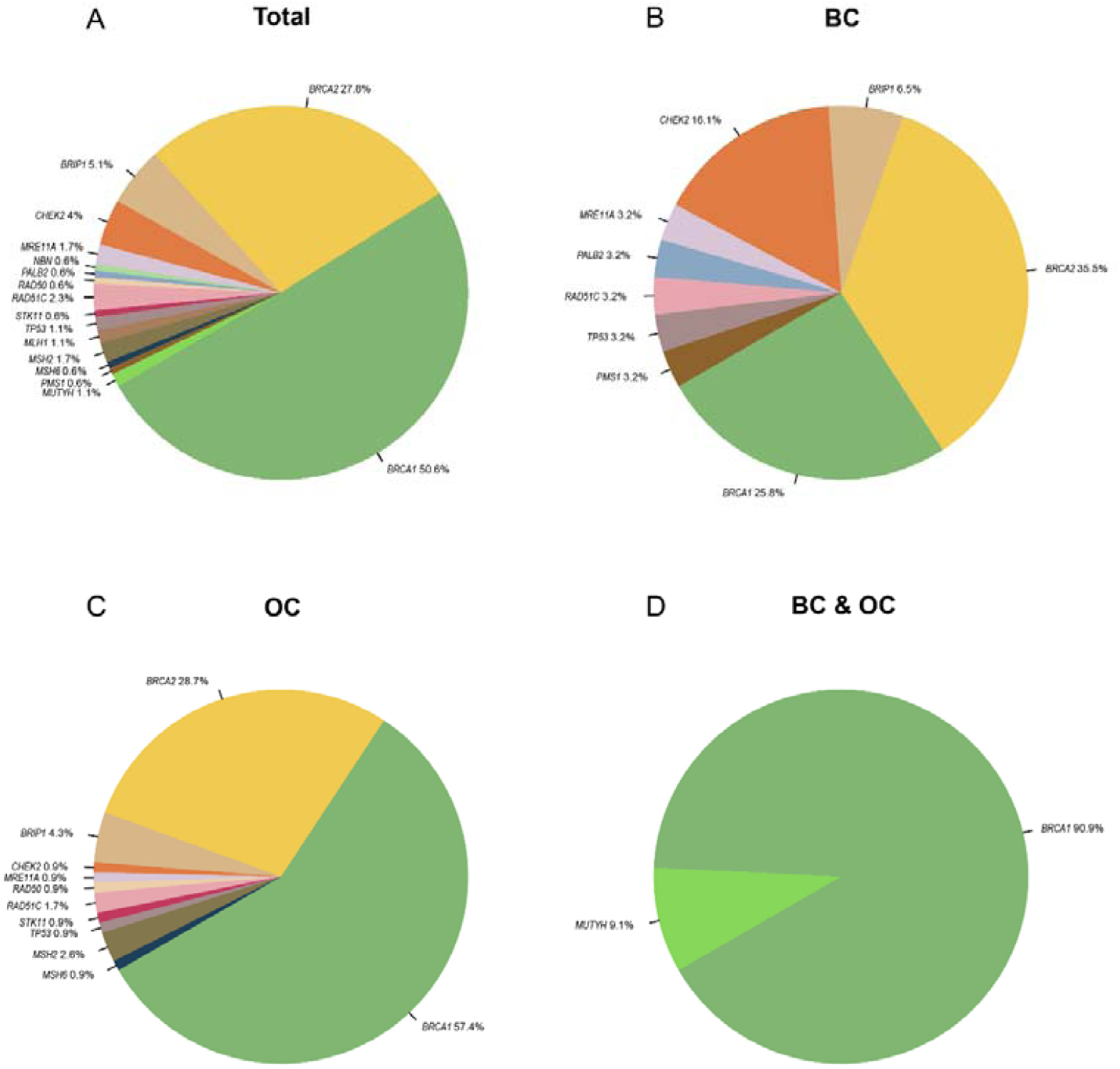
Deleterious mutation distribution in 21 HBOC susceptibility genes. (A) Overall 176 deleterious mutations were distributed in 21 genes among 172 patients of 882 individuals. Note that due to rounding, the sum is over 100%. (B) In 261 patients with breast cancer only, distribution of 31 deleterious mutations in 21 genes was identified. (C) In 429 patients with ovarian cancer only, distribution of 115 deleterious mutations in 21 genes was identified. (D) Distribution of 11 deleterious mutations distribution in 21 genes was identified in 19 patients with both breast and ovarian cancer.

**Figure 2.**
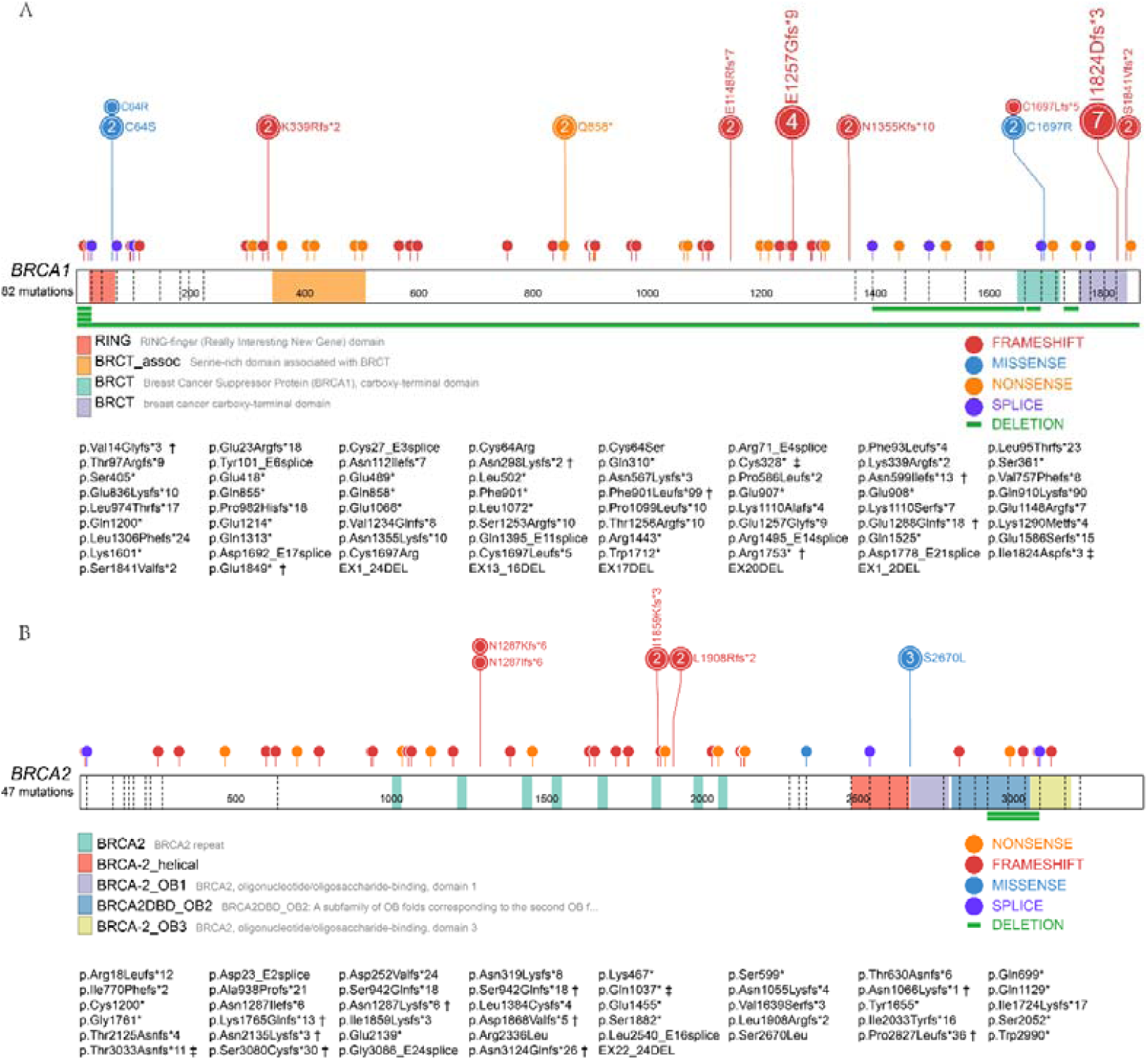
Spectrum of deleterious mutations detected in 882 individuals. (A) The spectrum of deleterious mutations in *BRCA1* gene. (B) The spectrum of deleterious mutations in *BRCA2* gene. Sticks represent mutation positions. The number represents the number of samples with the mutation (the unmarked represents 1). The green bars represent the deletion locations, and each segment represents a sample. The figure was made using ProteinPaint. † novel mutation; ‡ Chinese founder mutation

In the breast cancer only subgroup, 31 deleterious mutations were identified in 261 patients (Table 3). Most mutations occurred in *BRCA1* (66; 57.4%) and in *BRCA2* (33; 28.7%). An additional 16 mutations (13.9%) were found in 9 other susceptibility genes (Figure 1B). Deleterious *BRCA1* mutations consisted of 8 truncating (2 deletion, 2 frameshift, 2 nonsense, and 2 splice) mutations. Deleterious *BRCA2* mutations (2.7%) were 11 truncating mutations (9 frameshift, 2 nonsense). Among the other HR pathway genes, mutations were most commonly found in *CHEK2* (n = 5; 1.92%) and *BRIP1* (n = 2; 0.77%). In addition, mutations were also observed in *RAD51C, PALB2*, and *MRE11A* in 1 individual per gene. Only one Lynch syndrome gene mutation was identified in *PMS1* in the breast cancer subgroup. Among the other highly penetrant genes, mutations were found in *TP53* (n = 1; 0.38%) while no mutations were identified in *STK11, PTEN*, and *CDH1*.

In the ovarian cancer only subgroup (n = 429), 115 deleterious mutations were identified in 113 (26.3%) individuals (Table 3). Of these, 66 (57%) occurred in *BRCA1*, 33 (29%) in *BRCA2*, and 16 (14%) in 9 of 19 other susceptibility genes (Figure 1C). Deleterious *BRCA1* mutations consisted of 61 truncating (5 deletion, 35 frameshift, 16 nonsense and 5 splice) mutations and 5 known deleterious missense mutations. The 33 deleterious *BRCA2* mutations consisted of 30 truncating mutations (1 deletion,19 frameshift, 7 nonsense, and 3 splice mutations) and 3 known deleterious missense mutations. Among the homologous recombination (HR) pathway genes, the most frequent mutations were found in *BRIP1* (n = 5; 1.17%), and *RAD51C* (n = 2; 0.47%). Mutations in *CHEK2, MRE11A* and *RAD50* were identified in three individuals, respectively. For Lynch syndromes related genes (*MLH1, MSH2, MSH6, PMS1, PMS2*), deleterious mutations were identified in *MSH2* (n = 3) and *MSH6* (n = 1), accounting for 3.5% of all mutations in the ovarian cancer subgroup. Among the other highly penetrant genes, mutations were found in *TP53* (n = 1; 0.23%) and *STK11* (n = 1; 0.23%).

In the subgroup of subjects with disease histories of both breast and ovarian cancer (n = 19) (Table 3), a higher frequency of mutation rate was observed. Eleven (57.9%) subjects had a mutation, of which 10 had a mutation in *BRCA1*, and 1 had a *MUTYH* mutation (Figure 1D).

Further, 173 subjects do not have a cancer diagnosis but have a family history of cancer. In this subgroup, only 19 mutations were identified in 21 cancer susceptibility genes with a prevalence of 11.0% (Table 3), in which 10 were mutations in *BRCA1/2* genes, 2 in *BRIP1*, 2 in *MLH1*, 1 in *CHEK2*, 1 in *MRE11A*, 1 in *NBN*, 1 in *RAD51C*, and 1 in *MUTYH*. No mutations were found in *PALB2, RAD50, STK11, TP53, MSH2, MSH6*, and *PMS1* genes.

### Recurrent mutations, founder mutations, and novel mutations

In our cohort, recurrent mutations (n ≥ 3) were found in *BRCA1* p.Ile1824AspfsX3, *CHEK2* p.His371Tyr, *BRCA1* p.Glu1257GlyfsX9, and *BRCA2* p.Ser2670Leu (Table S1). And *BRCA1* p.Ile1824AspfsX3 was also one of Chinese founder mutations. The other Chinese founder mutations included *BRCA1* p.Cys328*, *BRCA2* p.Thr3033Asnfs*11, and *BRCA2* p.Gln1037Ter. No Ashkenazi Jewish or European founder mutations were observed. We confirmed 26 novel mutations that are not reported in public databases (ClinVar, UMD, LOVD, BIC) and literature. Of these, 7 in *BRCA1* (p.Val14Glyfs*3, p.Asn298LysfsX2, p.Asn599Ilefs*13, p.Phe901Leufs*99, p.Glu1288Glnfs*18, p.Arg1753Ter, p.Glu1849Ter), 9 in BRCA2 (p.Ser942Glnfs*18, p.Asn1066Lysfs*1, p.Asn1287LysfsX6, p.Lys1765Glnfs*13, p.Asp1868Valfs*5, p.Thr2125Asnfs*4, p.Pro2827Leufs*36, p.Ser3080CysfsX30, p.Asn3124Glnfs*26) (Figure 2), 3 in BRIP1 (p.Ser206Ter, p.Ser230Ter, p.Lys998AsnfsX5), 3 in RAD51C (p.Ser231Ter, p.Gln62Ter, p.Val41Glyfs*18), and 1 in CHEK2 (Leu303_E8splice), MSH2 (p.Asn412Metfs*22), NBN (p.Asn639Argfs*6), PMS1 (p.Tyr90*), respectively.

### Variants of uncertain significance (VUS)

The total VUS rate in our cohort was 38.55% (n = 340), and 406 VUS mutations (339 missense, 58 splice, 4 in-frame, 2 frameshift, 2 duplication, 1 deletion) were detected in 882 individuals. VUS mutations were most prevalent in *ATM* (n=53), followed by 29 VUS in *BRCA1* and 42 VUS in *BRCA2* (Figure 3). Apart from *BRCA1* and *BRCA2*, genes have insufficient interpretation of pathogenicity and benign polymorphism in the mutation classification, resulting in a high proportion of VUS. Of these mutations, VUS were found most frequently in *MRE11A* p.Met157Val (n = 8), *BRIP1* p.Gln944Glu (n = 8), and *ATM* p.His42Arg (n = 8). In addition, both *PMS1* p.Arg919Cys and *MSH6* p.Pro1082Ser were occurred in 6 individuals.

**Figure 3.**
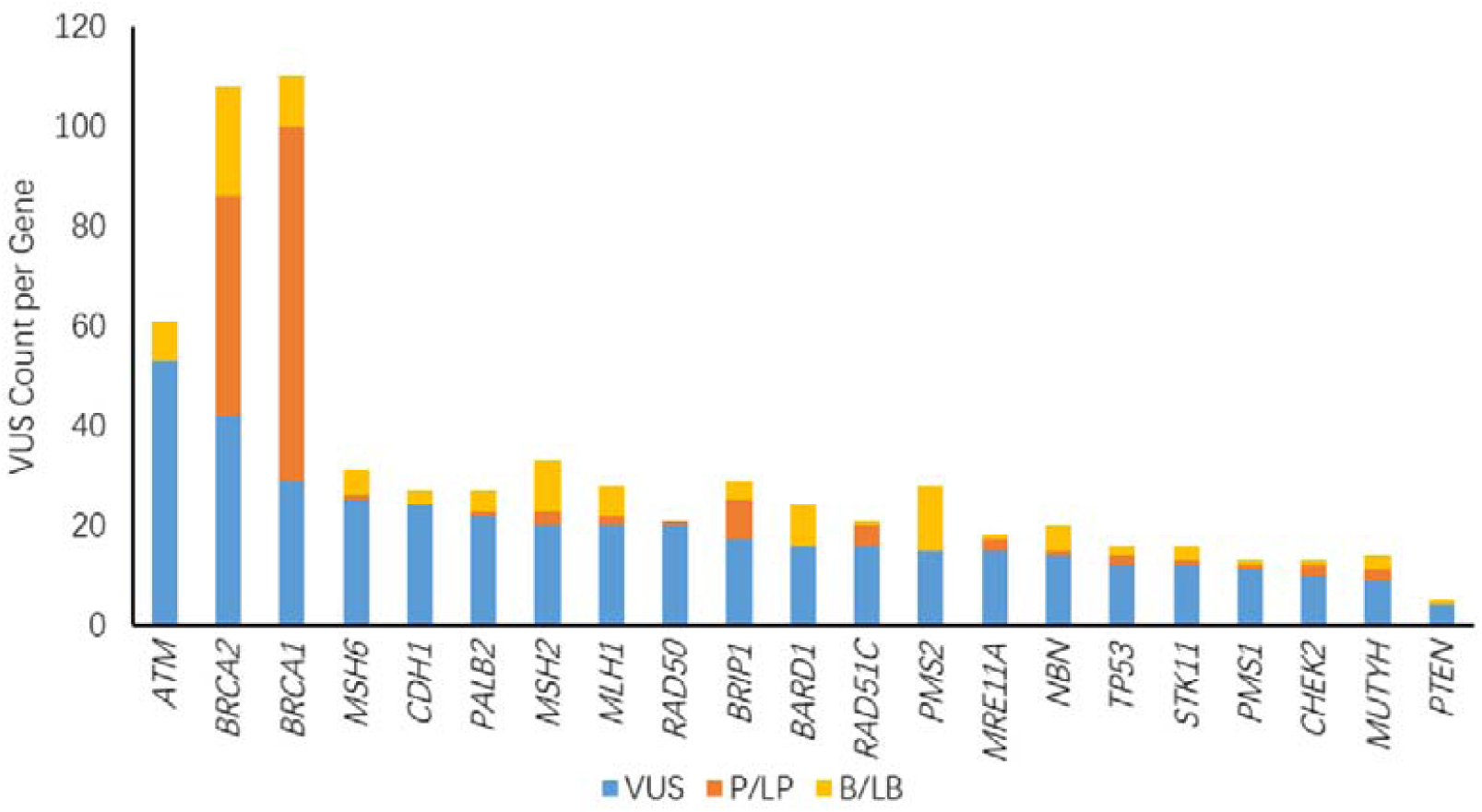
Overall proportion of VUS in 21 cancer susceptibility genes. VUS: variant of uncertain significance; P/LP: pathogenic or likely pathogenic; B/LB: benign or likely benign

### Mutation frequency in subgroup with different family histories and ages at diagnosis

The deleterious mutation rate for each subgroup according to age at diagnosis is detailed in Table 4. In the breast cancer only subgroup, the mean age at diagnosis was 39 among mutation positive probands and 40 among mutation negative probands, respectively (p = 0.66). In the ovarian cancer only cohort, the average age at diagnosis was 53 among positive probands and 54 among mutation negative probands (p = 0.90). In the breast and ovarian cancer cohorts, age at diagnosis was slightly older among mutation negative individuals compared to those positive for a mutation, however, the difference was not significant (p = 0.41) (Table 4). We also evaluated whether patient subjects with deleterious mutations in the 21 susceptibility genes were associated with a greater family history of breast and/or ovarian cancers than nonmutated patient subjects (Table 5). Among breast cancer patients, no significant association was identified between mutations and family history of either breast cancer or ovarian cancer. However, among ovarian cancer patients, individuals with mutations were more likely to have a family history of either breast or ovarian cancer (p < 0.05) (Table 5).

**Table 4.** Correlation between deleterious mutations and age in subgroup.

|  | **BC** |  | **OC** |  | **BC & OC** |  | **FHx** |  |
| --- | --- | --- | --- | --- | --- | --- | --- | --- |
| **Age (yr)** | Pos | Neg | Pos | Neg | Pos | Neg | Pos | Neg |
| **mean** | 39.7 | 39.8 | 53.6 | 53.4 | 54.8 | 60.3 | 41.1 | 40.4 |
| **median** | 39 | 40 | 53 | 54 | 52 | 59 | 35 | 39 |
| **range** | 26-5 | 20-62 | 27-7 | 24-7 | 44-6 | 39-8 | 27-71 | 13-7 |
| **p-value** | 0.6643 |  | 0.9033 |  | 0.4083 |  | 0.9027 |  |

Pos = positive, carried deleterious mutation
Neg = negative, no deleterious mutation

**Table 5.** Correlation between deleterious mutations and HBOC family history in subgroup.

|  | BC |  | OC |  |
| --- | --- | --- | --- | --- |
|  | Pos | Neg | Pos | Neg |
| **with HBOC family history** | 6 (2.3%) | 29 (11.1%) | 37 (8.6%) | 22 (5.1%) |
| **without HBOC family history** | 23 (8.8%) | 203 (77.8%) | 76 | 294 (68.5%) |
| *P*<sup>a</sup> | 0.246 |  | <0.05 |  |

a Chi-square test was used.

## Discussion

Using a HBOC multi-gene panel, we revealed prevalence of deleterious germline mutations among 882 subjects who were high-risk individuals and referred for Oseq-BRCA testing. This test utilizes liquid solution hybridization-based target enrichment and next generation sequencing to identify all types of variants in 21 HBOC genes. Our results support the views that the panel testing could increase the diagnostic detection rate of deleterious germline mutation compared with testing for *BRCA1/2* mutations alone. In our cohort, 172 (19.50%) subjects had a deleterious mutation, and 21.6% of deleterious mutations were in genes other than *BRCA1* and *BRCA2*. Our study is distinguished from other studies in the following ways. First, our large-size cohort is recruited from multiple-general hospitals across China which is a representative population for target population in China. The prevalence of mutations in this population was rarely reported in previous studies. In addition, our cohort was selected according to the NCCN guidelines, including breast cancer patients, ovarian cancer patients, and high-risk volunteers. Our results reflect the mutation frequency in individuals defined by the guidelines and have great clinical practical significance.

In the breast cancer only subgroup, the prevalence of *BRCA1* and *BRCA2* deleterious mutations were 3.07% and 4.21%, respectively. In a previous Chinese population-based study, Sun *et al*.^22^ reported that *BRCA1/2* deleterious mutations frequencies were 4.24% and 6.60% in early-onset breast cancer and familial breast cancer cohort, respectively, which is similar to our subgroup and those observed in other studies in China.^23,24^ However, the prevalence of *BRCA1/2* deleterious variants in breast cancer in other countries ranges from 9.3% to 18%.^25-27^ Among African Americans women, Churpek *et al*. reported that the prevalence in deleterious mutations in *BRCA1 and BRCA2* genes was 10% and 8%, respectively.^27^ Differences in the definition of early-onset or familial breast cancer and genetic testing methods for hereditary breast cancer between studies may influence results. Previous studies demonstrated that 4%-5% of breast cancer patients carried deleterious mutations beyond *BRCA1*/*2*, ^25-27^ which was consistent with our finding that 4.21% of this subgroup carried deleterious variants in neither *BRCA1* nor *BRCA2*. The third commonly mutated gene was *CHEK2* in our study, which encodes a checkpoint kinase 2 interacting with cell cycle regulators and DNA repair proteins. And the deleterious mutation of *CHEK2* would increase the risk of breast cancer.^14^ Five patients carried *CHEK2* deleterious mutations, 4 in p.His371Tyr and 1 in c.908+2T>A. Although the recurrent mutation p.His371Tyr in *CHEK2* was marked as variant uncertain significance in ClinVar database, we interpreted it as likely pathogenic variants. This mutation results in the change of a Histidine to a Tyrosine at position 371 of the CHEK2-encoded protein. Baloch *et al*. found this mutation occurred in a domain of protein kinase activity that plays an important role in DNA damage repair.^28^ This mutation is a suspected disease-causing mutation with 1 strong pathogenicity (PV3: functional studies supportive of a damaging effect) (6) and 1 moderate pathogenicity (PM2 low frequency in 1000 Genomes Project). Noteworthy, our study found only 1 breast cancer patient carried *PALB2* mutation but no patients carried ATM mutation, while *ATM* and *PALB2* mutation were commonly identified in other studies.^25-27^

The frequency of *BRCA1/2* mutations was 23.07% in the ovarian cancer only subgroup, 15.38% for *BRCA1* and 7.69% for *BRCA2*, respectively. The mutation rate of *BRCA1/2* in our ovarian cancer subgroup was slightly higher than that in other studies that found *BRCA1/2* deleterious mutations rated from 13% to 15%.^29-31^ Overall, *BRCA1/2* mutation accounted for 86% of total mutations in hereditary ovarian cancer, and the *BRCA1* mutation rate was more pronounced than the *BRCA2* in ovarian cancer patients, which is similar to the results from both Ang Li *et al*. ^32^ (2018, n = 1331, *BRCA1* for 17.1% and *BRCA2* for 5.3%) and Norquist *et al*. ^29^ (2016, n = 1915, *BRCA1* for 9.5% and *BRCA2* for 5.1%). Apart from B*RCA1/2* mutation, 0.9% of the subgroup carried *BRIP1* mutation (BRCA1-interacting protein C-terminal helicase 1), which is comparable to other studies in which the prevalence ranges from 0.8% to 1.5%.^29,33^ BRIP1, a member of the BRCA-Fanconi anemia DNA repair pathway, is one of ovarian cancer moderate-risk genes and *BRIP1* mutations is associated with a 10%-15% increased risk of lifetime ovarian cancer.^34^ Reviewing 5 patients with *BRIP1* deleterious mutations, all subjects had a family history of cancer (ovarian cancer, breast cancer, pancreatic cancer, colon cancer, gallbladder cancer). This data suggests that *BRIP1* mutation may be the pathogenic cause in ovarian cancer patients with a family history of cancer. In reviewing the mutations in mismatch repair genes (MMR; *MLH1, MSH2, MSH6, PMS2*), mainly causing Lynch syndrome, were of low frequency in our subgroup (n = 4; 0.93%). However, in our cohort, MMR mutations only occurred on the *MSH2* and *MSH6* genes, and no mutations in *MLH1* were found, which is different from the spectrum of hereditary colorectal cancer. This phenomenon also occurred in the study of Norquist *et al*. (7 of 8 MMR mutations occurred in *PMS2* or *MSH6*).^29^ Although the values of these genes are unknown with respect to risk assessment, we cannot completely rule out benefit of these genes when doing genetic testing in ovarian cancer.

In the subgroup of subjects with a diagnosis of both breast and ovarian cancer, high deleterious mutation rates (52.63% and 5.26%) were observed in *BRCA1* and *MUTYH*. Kwong *et al*. (2018, n = 20) reported that the prevalence of mutation Chinese patients with breast cancer complicated with ovarian cancer were 40% and 20% for *BRCA1* and *BRCA2*, respectively.^35^ Walsh *et al*. reported that the frequencies *of BRCA1* and *BRCA2* mutation were 38.71% and 22.58%, and there were three additional subjects carried *BRIP1, CHEK2*, and *MRE11A* mutations, respectively.^30^ It seems that the frequency of *BRCA1* mutations in this subgroup is higher than that of carrying *BRCA2* mutation, although the prevalence of a diagnosis of both breast and ovarian cancer is relatively low in domestic and international research. Given limited sample size, more evidence is needed to support this point.

The rate of VUS in a similar multi-gene panel study (27 genes) was 32.7%, in which they use a panel with fewer genes but with *BRCA1/2* included.^36^ Indeed, VUS rate in our cohort was 38.55%, which is slightly higher than the results of previous study. It is possible that the incidence of breast cancer and ovarian cancer in the Chinese population is lower than that in Caucasians, and the variants are relatively sporadic.

*ATM* has the most frequent VUS detected due to the long transcript length. When we exclude the length of CDS to make comparisons, it shows that *RAD51C* has the greatest number of VUS in per 1000 base, up to 14.15 (Table S2). According to the NCCN guidelines, *RAD51C* specifically increases the risk of ovarian cancer. In our ovarian cancer patients, only one deleterious mutation of *RAD51C* was detected, which is relatively low compared with previous populations.^37^ This may due to the lack of reports on *RAD51C* mutation in Chinese population.

Identification of a deleterious variants in a cancer susceptibility gene allows identification of eligible patients for surveillance screening, and it may provide targeted therapy and prevention strategies for both patients and family members. Clinical interventions and recommendations of *BRCA1* and *BRCA2* mutation carriers have been well established and widely used in clinical practice. Most genes in our panels (*CDH1, MSH2, MLH1, MSH6, PMS2, PTEN, STK11*, and *TP53*) had corresponding current management suggestions in the NCCN guidelines. However, other moderate penetrance genes (*BARD1, RAD50, ATM, BRIP1, CHEK2, NBN, PALB2, RAD51C*) are not available in the management guidelines, while mutation in these genes were found in 2.60% subjects. When encountering these mutations, it is a big challenge for clinicians. It is necessary to combine the family history and personal history to make a medical decision. Therefore, guidelines recommend that multi-gene testing is ideal in the context of professional genetic expertise for pre- and post-test counseling.

In conclusion, we reported the successful utility of multiple gene testing for identification of HBOC relevant risk gene mutations in a large-scale mutation screening. This is the first clinical investigation of the mutation spectrum with a multiple gene panel among high-risk Chinese individuals with a suspected HBOC risk. Results of this study indicated that multi-gene panel testing can identify more individuals with relevant cancer risk gene mutations than *BRCA1/2* genetic testing alone. Although current NCCN guidelines recommend the management of patients with mutations in majority of risk genes, the clinicians should be prepared to deal with the VUS and mutations in moderate penetrance genes. Our findings provide insights for the clinician to consider multi-gene tests to diagnose cancer predisposition in clinical practice.

## Abbreviations

HBOC: hereditary breast and ovarian cancer
PARP: poly-ADP-ribose polymerase
PCR: polymerase chain reaction
DGGE: denaturing gradient gel electrophoresis
NCCN: National Comprehensive Cancer Network
TRFs: test requisition forms
InDels: insertions and deletions
GATK: Genome Analysis Toolkit
CNVs: copy number variants
HGVS: Human Genome Variation Society
ACMG: American College of Medical Genetics
VUS: variant of uncertain significance
HR: homologous recombination
FA: Fanconi anaemia

## Acknowledgments

This work was supported by National Key R&D Program of China (2016YFC0905400), Guangzhou Science and Technology Plan Projects (Health Medical Collaborative Innovation Program of Guangzhou, 201400000004-5 and 201803040019) and Special Foundation for High-level Talents of Guangdong (2016TX03R171).

## Supplementary Data

Supplement Table S1 List of deleterious mutations

Supplement Table S2 Number of VUS mutations per kb in 21 genes

Supplement Figure S1 Distribution of *BRCA1* and *BRCA2* deleterious mutations

