## Supplementary figures and images for "Frequency of mutations in 21 hereditary breast and ovarian cancer susceptibility genes among high-risk Chinese individuals"

### Supp. Figure S1

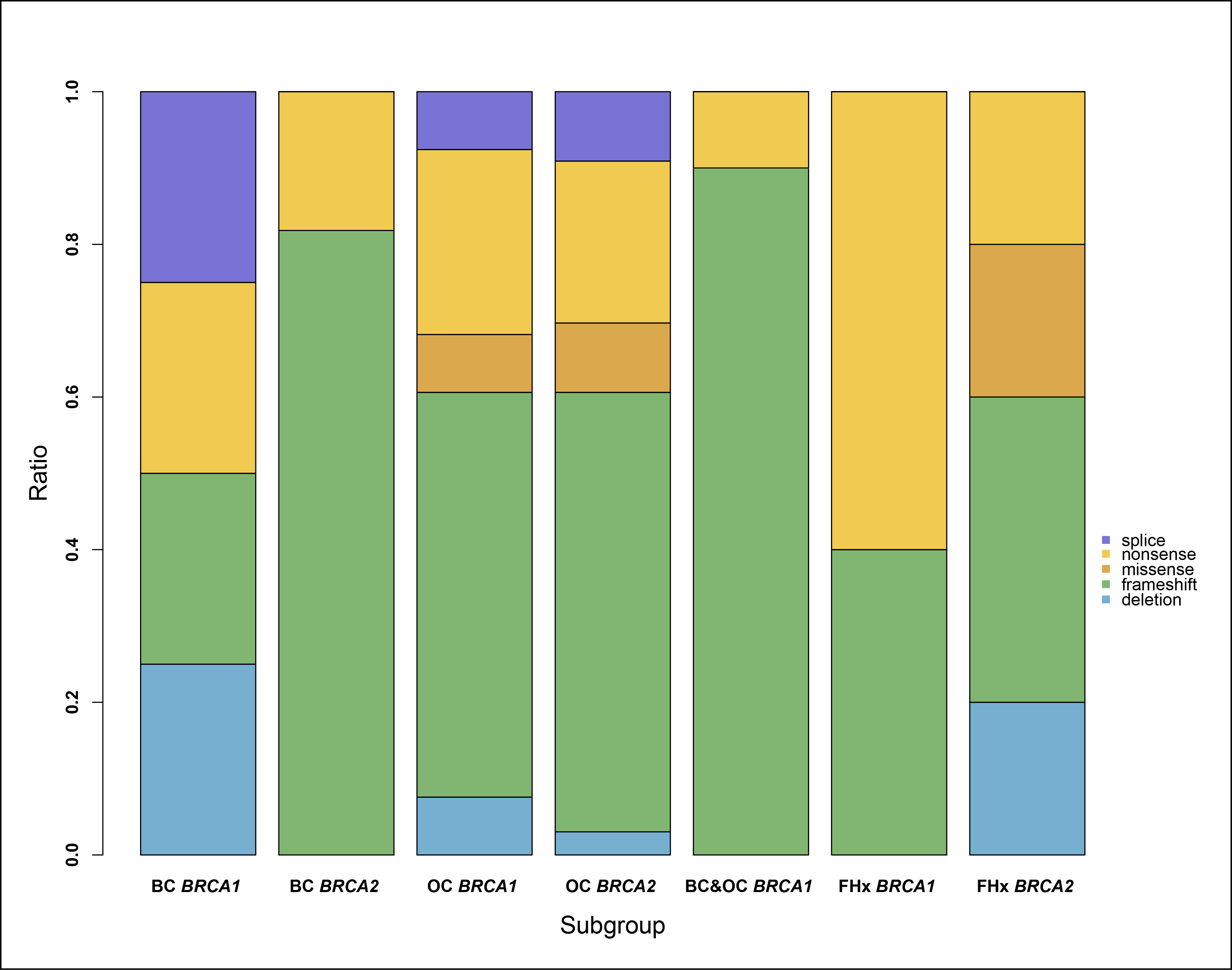
